# Amyloid-Induced Network Resilience and Collapse in Alzheimer’s Disease: Insights from Computational Modeling

**DOI:** 10.64898/2026.02.09.704344

**Authors:** Ediline L. F. Nguessap, Damien Depannemaecker, Fernando F. Ferreira

## Abstract

Alzheimer’s disease (AD) is marked by progressive synaptic loss and cognitive decline, with amyloid-beta ( *A β* ) accumulation playing a central pathological role. While the molecular effects of *A β* are well-characterized, the cascade from local synaptic dysfunction to largescale network collapse remains poorly understood. We present a biophysically grounded computational model that integrates small-world network topology, Izhikevich neuron dynamics, short-term plasticity (STP), and A *β*-induced synaptic pruning to investigate how amyloid burden degrades functional brain connectivity over time. Crucially, we simulate the evolving network dynamics across disease stages, revealing how firing rate, synchrony, variability (CV), and Fano factor evolve in response to progressive structural degradation. Our results demonstrate a two-phase deterioration: initial compensatory dynamics followed by abrupt disintegration of coordinated activity. This temporal dissociation mirrors clinical observations and offers mechanistic insight into AD progression. We further identify a critical amyloid threshold beyond which network collapse becomes inevitable, preceded by early-warning indicators such as declining global efficiency. Network topology, synaptic time constants, and repair mechanisms strongly influence resilience, with small-world networks showing delayed collapse and greater functional compensation. Finite-size scaling reveals that larger networks experience earlier and sharper phase transitions.These findings unify molecular, structural, and dynamical perspectives, suggesting that fluctuations in global efficiency and synchrony could serve as sensitive biomarkers for early detection. Our model offers a predictive, multiscale framework for understanding AD progression and evaluating therapeutic strategies.

## 1 Introduction

Alzheimer’s disease (AD) is the most common form of dementia and is characterized by progressive cognitive decline, synaptic loss, and neuronal death. A hallmark of AD pathology is the accumulation of amyloid-beta ( *A β* ) plaques, which disrupt synaptic function and trigger a cascade of neurodegenerative processes [1,2]. Despite extensive experimental studies, the link between molecular-scale *A β* accumulation and large-scale network collapse remains poorly understood. Clinical observations suggest that functional brain networks exhibit altered connectivity and reduced efficiency long before the appearance of overt structural damage or cognitive symptoms [3,4].

Computational modeling has emerged as a powerful approach to explore the mechanisms of disease progression across spatial and temporal scales. Previous work has shown that AD-related damage propagates non-randomly across the brain, preferentially targeting hub regions and disrupting integration [5,6]. Other studies have explored the impact of excitability, inflammation, or neurovascular coupling on network vulnerability [7,8], and some have incorporated spreading models of pathology over structural connectomes. However, many of these models focus solely on structural disconnection or static analysis, and few address how synaptic failure translates into dynamic alterations in spiking activity.

Moreover, experimental evidence indicates that synaptic plasticity and compensatory sprouting are active during the early stages of AD [9], suggesting that the brain attempts to counteract degradation before reaching a critical threshold. Understanding how these compensatory mechanisms interact with network architecture and neuronal dynamics is essential for identifying early biomarkers and intervention windows.

While prior studies have modeled amyloid-driven synaptic degradation using structural networks [5, 6], they often omit dynamic spiking activity, synaptic plasticity, or repair mechanisms, limiting their ability to capture the full trajectory of Alzheimer’s progression. Other models have addressed hyperexcitability [7] or propagation dynamics [8], but typically neglect functional metrics or do not incorporate biologically grounded thresholds for collapse. Moreover, few frameworks integrate both network topology and intrinsic neuronal excitability when evaluating resilience. Additionally, many existing models rely on rate-based or abstracted node dynamics, which are less suited to capturing temporal patterns such as spike bursts, firing variability which are features increasingly linked to early AD signatures in electrophysiological data [10]. A spiking-neuron-based framework is therefore essential for capturing dynamic signatures of collapse and compensatory coordination.

In contrast, our work presents a unified modeling framework that incorporates small-world topology, Izhikevich neuronal dynamics, short-term synaptic plasticity (STP), and A *β*-induced synaptic pruning, along with activity-dependent repair. Short-term plasticity (STP), in particular, modulates synaptic efficacy on the timescale of milliseconds to seconds, shaping the propagation of activity in networks undergoing stress. Its role in early compensation has been observed experimentally, but is rarely incorporated into computational

AD models. This enables us to capture both structural degradation and emergent functional collapse, characterize early-warning signals, and explore how both synaptic timescales and neuronal excitability influence the temporal dynamics of network failure.

In the following sections, we develop and analyze this biophysically motivated network model to simulate the progressive effects of *A β* toxicity on neural connectivity and dynamics. We identify critical thresholds, quantify early indicators of collapse, and assess how network topology, plasticity, and intrinsic dynamics shape the trajectory of disease progression.

## 2 Models

### 2.1 Network Construction

The neuronal network was implemented as a directed Watts-Strogatz small-world network with *N* = 10^4^ neurons and average degree *k* = 20, parameters chosen to match the connectivity statistics observed in mammalian cortical microcircuits [11]. All synaptic weights were initialized to *W* _0_ = 1, representing normalized connection strengths. This uniform initialization allows clear observation of amyloid-induced degradation effects without confounding by initial weight heterogeneity. Neurons were removed from the network when their incoming synaptic weights fell below *W* _0_ / ( 2 *e* ), a threshold derived from the minimum functional connectivity required for signal propagation in spiking network models [12]. This removal condition models synapse elimination when postsynaptic potentials drop below 18 % of initial strength, consistent with electrophysiological measurements in AD pathology [13].

### 2.2 Amyloid Dynamics

Amyloid accumulation followed logistic growth with microglial clearance:

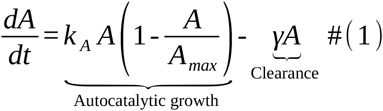

with *k* _*A*_ = 0.03 month ^-1^ producing the characteristic 15-20 year progression from preclinical to dementia stages [14], *A*_max_ = 150 CL (Centiloid units), and *γ* = 0.005 month ^-1^ models microglial activity [15]. When amyloid levels exceeded the toxicity threshold *A*_th_ = 25 CL which matches PET-defined amyloid positivity [16], Synaptic weights decayed exponentially as follows: 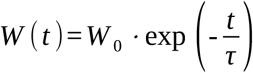 with pruning probability defined as :

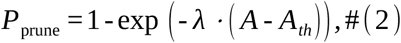

The toxicity factor *λ* in *CL*^-1^ was derived from post-mortem studies correlating amyloid load with synapse loss [17]. Surviving synapses weaken exponentially with time constants *τ ∈* [ 60,200 ] months matching the timescale of cognitive decline in prodromal AD [14].

In our model, the synapse decay time constant ( *τ* _synapse_ ) governs the rate of synaptic weakening after amyloid levels cross a pathogenic threshold ( *A*_*th*_ ). While *τ* _*synapse*_ and *A*_*th*_ are mathematically independent, their interplay determines the transition from amyloid accumulation to network failure, mirroring the delayed synaptic degeneration observed in Alzheimer’s disease (AD) [13,17]. The model employs a quasi-static approximation justified by distinct biological timescales:

- Slow pathological changes: Amyloid accumulation ( *τ* _pathology_ in years) and synaptic degradation ( *τ* _*s*_ ∼ 5 - 20 years) [18, 19].
- Fast neuronal dynamics: Action potentials ( *τ* _spiking_ in milliseconds) and STP ( *τ* _facil_ ∼ 50 - 1000 ms) [20,21]

This separation allows static network snapshots (pre-collapse, collapse-onset, post-collapse) to be simulated independently while preserving the cascade from molecular dysfunction to network failure [5]. The approximation is validated by Clinical observations of amyloid deposition preceding cognitive decline by ∼ 10 years [3] and Computational studies confirming timescale decoupling in neurodegenerative diseases [22].

### 2.3 Repair and reconnection mechanism

The model incorporates two biologically observed compensation mechanisms: synaptic repair and axonal sprouting. Existing synapses repair with probability:

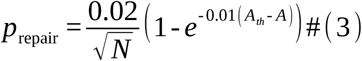

Each existing synapse ( *i, j* ) is strengthened with probability *p*_repair_ as:

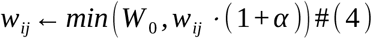

*α* = 3 % is the repair strength factor matching measured LTP magnitudes in early AD [1]. New connections form at half the repair rate probability 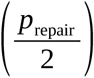 with initial weight 0.5 ⋅ *W* _0_, simulating axonal sprouting [13].

### 2.4 Neuronal Dynamics with Short-Term Plasticity (STP)

We extracted adjacency matrices at critical stages: (1) pre-collapse (healthy state), (2) collapse onset (amyloid threshold), and (3) post-collapse (disintegrated state). Neuronal dynamics were simulated on each static snapshot using the Izhikevich model to isolate the impact of connectivity changes on spiking activity. While this assumes synaptic weights remain fixed during simulation windows, this quasi-static approximation is justified by the timescale separation between amyloid accumulation in months and years [19, 23] and spiking dynamics in milliseconds [11,21]. This timescale separation is supported by computational studies of AD [22,24]. To study the impact of synaptic dynamics on neuronal activity, we simulate the Izhikevich neuron model coupled with Short-Term Plasticity (STP). The neuronal and synaptic dynamics are governed by the following equations:

#### a) Neuron Dynamics

The membrane potential *v*_*i*_ and recovery variable *u*_*i*_ for each neuron evolve according to the Izhikevich model [21]:

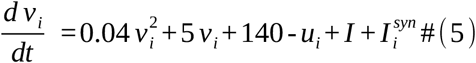

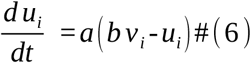

with spike resetting:

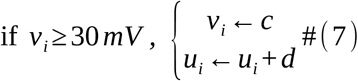

We explore variations in intrinsic excitability by adjusting the parameters *a* and *d*, which modulate neuronal recovery and after-spike reset, respectively.

#### b) Short-Term Plasticity (STP) Dynamics and input current

Short-term synaptic plasticity (STP) is implemented as a dynamic modulation of synaptic efficacy, governed by a recovery time constant and usage parameter.

For each synapse ( *i*→ *j* ), we define: *x*_*ij*_ ( *t* ) the facilitation (use of synaptic efficacy); *R*_*ij*_ ( *t* ) : the fraction of synaptic resources available; *U* : baseline utilization; *τ* _*facil*_ and *τ* _*rec*_ the time constants for facilitation and recovery.

After a spike from neuron:

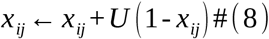

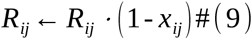

Between spikes, the variables evolve as:

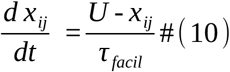

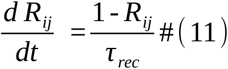

The effective synaptic weight becomes:

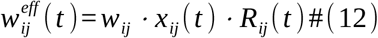

And the synaptic input to neuron *j* is:

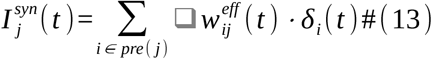

where *δ*_*i*_( *t* ) equals 1 if neuron *i* spikes at time *t*, and 0 otherwise. In our computational framework, the implementation of repair and reconnection mechanisms is biologically motivated and temporally aligned with amyloid-beta ( *Aβ* ) progression expressed in Centiloid ( *CL* ) units. Research suggests that ( *Aβ* ) begins to accumulate silently during the preclinical phase of Alzheimer’s disease, with neurotoxic effects becoming detectable beyond 25 CL which is the threshold for ( *Aβ* )-positivity [25]. Our model assumes that neuronal dysfunction may begin subtly in this early phase, prompting compensatory repair mechanisms such as axonal sprouting, synaptic reformation, or homeostatic plasticity [1,26].

Experimental data suggest that once ( *Aβ* ) levels surpass the threshold and continue to rise, especially as they approach 75CL (associated with cognitive impairment), the efficiency of these endogenous repair mechanisms diminishes [2,27]. However, our model incorporates pre-symptomatic repair mechanisms that become inactive when amyloid levels ( *A* ) reach the positivity threshold ( *A* ≥ 25 *CL* ).

The network equations are solved numerically using the 4th-order Runge-Kutta method to ensure accurate integration of neuronal and synaptic dynamics with time step of 1 *ms*.

### 2.5 Graph Metrics

We compute the following graph metrics to quantify network integrity:

#### 2.5.1 Largest Strongly Connected Component (LSCC)

The LSCC is the largest subset of neurons in which every neuron is reachable from every other neuron in the subset via directed paths. The size of the LSCC is normalized by the total number of neurons : 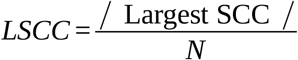, where |Largest SCC| is the number of neurons in the largest strongly connected component.

#### 2.5.2 Global Efficiency

Global efficiency measures the efficiency of information transfer across the network. It is defined as the average inverse shortest path length between all pairs of neurons: 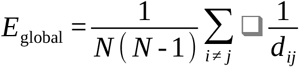, where *N* : Total number of neurons and *d*_*ij*_ : Shortest path length between neurons *i* and *j*.

#### 2.5.3 Susceptibility of the LSCC/ Fluctuation of the mean of the LSCC

To identify critical points in the network, we calculate the susceptibility *V* of the LSCC, which measures the fluctuations of the LSCC in relation to its average value in several independent projects.

The susceptibility is defined as: 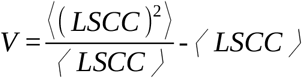, where: (⟨ *LSCC* ⟩ is the average of the LSCC in *n*_real_ activities: 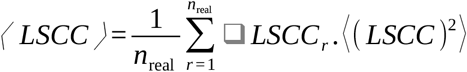 is the mean square of the LSCC in *n*_real_ realizations: 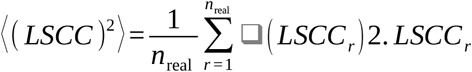 is the value of LSCC the *r*-th realisation.

This susceptibility measure evolves dynamically, identifying time-dependent critical points in the network; notably the critical time and critical amyloid level where connectivity collapses. Fluctuations in *V* ( *t* ) reflect temporal instability in LSCC size, signaling progressive network deterioration or phase transitions as the system approaches collapse.

## 3 Structural Degradation and Network Resilience

### 3.1 Network Degradation: LSCC Collapse and Critical Threshold

The constructed network is of average clustering coefficient of ≈ 0.5 and the shortest path length of ≈ 4.4. Figure 1 provides a comprehensive illustration of the temporal evolution of Alzheimer’s pathology and its impact on network connectivity and compensatory mechanisms. Panel (a) shows the sigmoidal progression of amyloid-beta accumulation, modeled through logistic growth dynamics with three phases: slow buildup ( = 0 - 150 months, *R*^2^ = 0.98 ), rapid expansion ( *t* = 150 - 300 months), and saturation (t > 300 months). Two key thresholds are marked: the toxicity threshold at 25 CL (dash red), where synaptic degradation begins, and the collapse threshold at ≈ 75 CL(solid red), corresponding to irreversible network failure. This growth pattern mirrors clinical observations, highlighting a long silent phase, a rapid expansion stage, and eventual saturation.

**Figure 1.**
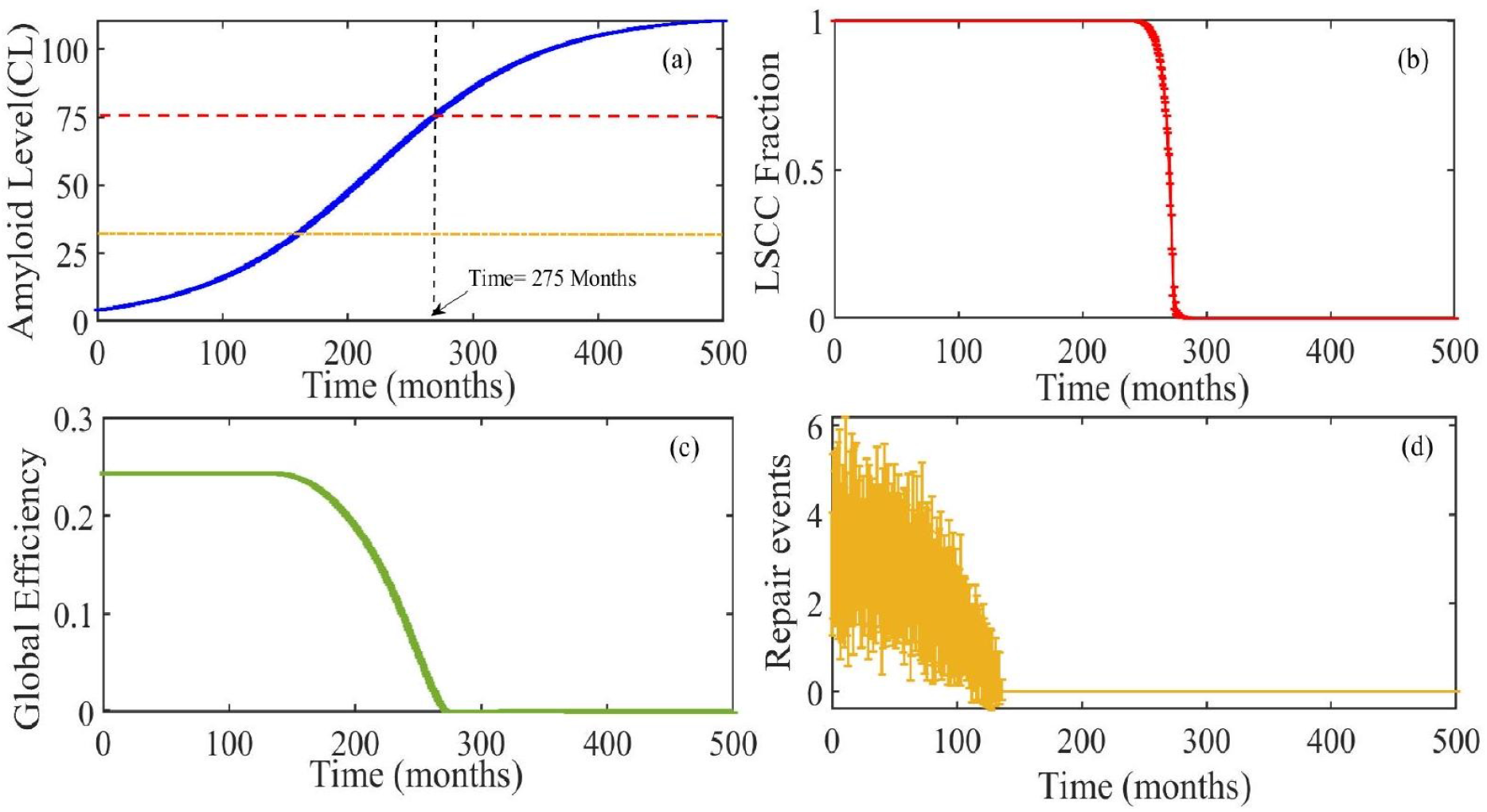
Network degradation under progressive amyloid-beta accumulation: (a) *A β* accumulation over time showing two key thresholds: 25 CL (toxicity onset) and 75 CL (collapse). (b) Fraction of the largest strongly connected component (LSCC) collapses after 75 CL. (c) Global efficiency (GE) declines earlier, indicating functional degradation before structural collapse. (d) Repair events peak early and cease once *A β* exceeds 25 CL Parameters: *τ* = 120 months, *N* = 10^4^. Error bands: SEM ( *n* = 20 ).

Panel (b) captures the fraction of the Largest Strongly Connected Component (LSCC), a structural marker of global network integration. The LSCC remains largely intact during early amyloid buildup but collapses abruptly once *A β* surpasses the 75 CL threshold. This sharp, nonlinear decline indicates a critical transition resembling a phase shift, where resilience gives way to fragmentation.

In contrast, panel (c) shows that global efficiency (GE), a measure of functional integration, begins to decline significantly before LSCC breakdown. This early decline acts as a leading indicator of structural collapse, consistent with the theory of critical slowing down, and suggests that functional disconnection may serve as a sensitive biomarker of preclinical Alzheimer’s progression.

Finally, panel (d) illustrates the trajectory of repair events, such as synaptic reinforcement and axonal sprouting. These compensatory processes peak just below the 25 CL threshold and cease rapidly thereafter, indicating a narrow window of effective plasticity during early amyloid accumulation. This dynamic justifies the model’s assumption that repair mechanisms operate only below the toxicity threshold.

Collectively, the figure encapsulates the multi-phase degradation of brain networks in Alzheimer’s disease, linking molecular pathology to the functional and structural collapse of neuronal connectivity.

### 3.2 Effects of Network Size and Finite-Size Scaling

The structural vulnerability of neuronal networks under amyloid stress varies and depend on the network size. To further investigate it, we systematically analyzed the effect of network size on the timing and nature of connectivity collapse Figure 2 examines how network size influences the progression and abruptness of connectivity loss under amyloid-induced degeneration, revealing a counterintuitive finding: larger networks exhibit earlier and more abrupt collapse. Panel (a) displays the evolution of the fraction of the Largest

**Figure 2.**
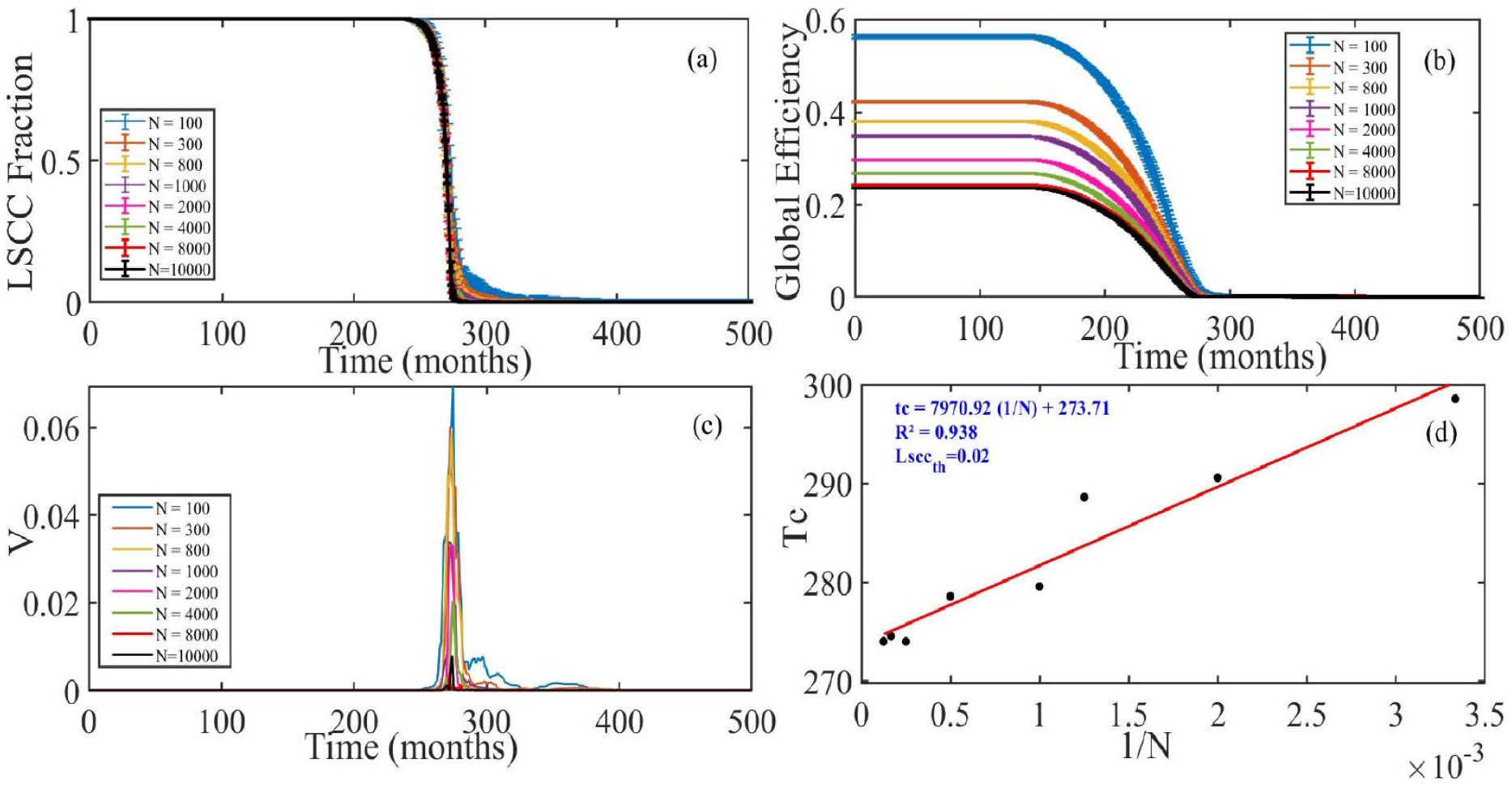
Network size modulates collapse timing and sharpness: (a) LSCC fraction over time for networks with *N* = 100 to 10000. Larger networks collapse earlier and more abruptly. (b) GE declines sooner in larger networks. (c) LSCC susceptibility *V* peaks near the collapse point and sharpens with increasing*N* . (d) Collapse time *t*_*c*_ scales inversely with *N*, indicating a finite-size transition. Error bands: SEM ( *n* = 20 ).

Strongly Connected Component (LSCC) over time for different network sizes ( = 100 to 10000 ). Contrary to expectations that increased size might enhance resilience, the results show that LSCC fragmentation occurs sooner in larger networks. The inset highlights that after collapse, the LSCC remains below 0.1 for larger networks, indicating a sharper and more permanent breakdown of global connectivity. Panel (b) confirms this vulnerability by showing global efficiency (GE) as a function of time: networks with more neurons suffer earlier declines in functional efficiency. This result implies that while larger networks have higher baseline integration, they are also more sensitive to perturbations once degradation begins.

Panel (c) introduces the susceptibility *V* of the LSCC, which measures fluctuations across realizations and peaks near the transition point for all network sizes. As *N* increases, this peak becomes sharper and shifts leftward, consistent with the hallmark of a second-order phase transition and supporting the notion of critical slowing down in AD progression [28]. Finally, panel (d) presents a finite-size scaling analysis, plotting the critical collapse time *t*_*c*_ (defined as the time at which LSCC drops below 0.02 ) against 1 / *N* . The linear relationship observed ( *R*^2^ = 0.938 ) suggests that the critical point becomes increasingly well-defined as the system size grows, aligning with finite-size scaling theory in statistical physics.

Together, these findings indicate that larger neuronal networks, despite their higher initial robustness, are more prone to sudden and earlier transitions into dysfunctional states under amyloid stress. This aligns with prior computational work by Kashyap et al. [6], which reported more abrupt transitions in large-scale systems due to amplified sensitivity to structural perturbations. These results underscore the importance of accounting for network size in models of neurodegeneration and suggest that individual variability in connectome scale may partly explain differing disease trajectories in Alzheimer’s patients.

### 3.3 Effects of Synaptic Time Constant (*τ* _*s*_ )

The differential effects of *τ* _*s*_ values reveal a fundamental trade-off between synaptic resilience and network stability in Alzheimer’s disease (AD) progression. Shorter synaptic time constants ( *τ* _*s*_ ) accelerate network degradation, while longer *τ* _*s*_ delays the collapse of the LSCC.

To explore how the temporal properties of synaptic weakening influence network resilience, we analyzed the effect of varying the synaptic decay time constant *τ* _*s*_ on network degradation dynamics. Figure 3 presents the evolution of both the Largest Strongly Connected Component (LSCC) and global efficiency (GE) for networks subjected to different values of *τ* _*s*_. These simulations reveal a critical trade-off between synaptic vulnerability and the timing of network collapse. In Panel (a), networks with shorter decay constants (e.g., *τ* _*s*_ = 60 months) exhibit early and rapid LSCC collapse, indicating accelerated structural degradation. This dynamic corresponds to more aggressive forms of Alzheimer’s disease, where synaptic integrity deteriorates swiftly and irreversibly [29].

**Figure 3.**
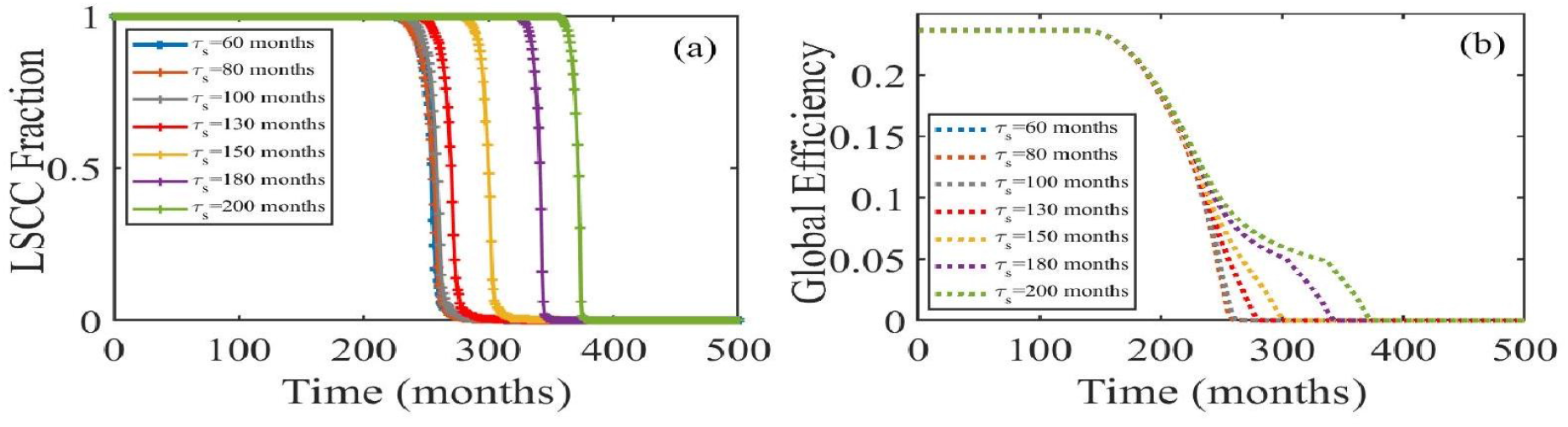
Effect of synaptic decay time *τ* _*s*_ on network resilience: (a) LSCC trajectories for different *τ* _*s*_. Larger *τ* _*s*_ delays collapse. (b) GE declines earlier for shorter *τ* _*s*_. Networks with slower synaptic decay retain functional connectivity longer.

Conversely, longer decay constants (e.g., *τ* = 200 months) delay the collapse, preserving the LSCC even as amyloid levels rise. This resilience reflects a compensatory phase in which network topology buffers against toxicity; an effect analogous to the cognitive reserve observed in some AD patients [30]. Panel (b) further illustrates that GE deteriorates significantly earlier in fast-decay scenarios, emphasizing that functional disintegration precedes structural breakdown. Notably, the time lag between amyloid crossing the toxicity threshold ( *A*_*th*_ = 25 CL) and LSCC failure increases with *τ* _*s*_, indicating a biphasic progression: an initial period of subclinical compensation, followed by an abrupt transition to system-wide failure.

This dissociation of functional and structural collapse highlights the importance of *τ* _*s*_ as a tunable parameter that captures individual differences in disease trajectories. Moreover, the results suggest that patients with longer synaptic decay times may benefit more from early therapeutic interventions aimed at preserving synaptic efficacy. These findings are consistent with clinical studies showing that synaptic biomarkers decline years before cognitive symptoms appear [4,31]. By operationalizing *τ* _*s*_ within a dynamic framework, our model links microscale degeneration to macroscale network failure, providing mechanistic insight into how subtle changes in synaptic resilience modulate Alzheimer’s progression.

To further quantify the impact of synaptic decay dynamics on network resilience, we investigated the relationship between the synaptic time constant *τ* _*s*_ and the critical collapse time *t*_*c*_, defined as the time at which the LSCC drops below 0.02 . This analysis is presented in Figure 4, which shows a clear monotonic relationship: as *τ* _*s*_ increases, the value of *t*_*c*_ is systematically delayed. This trend demonstrates that longerlasting synapses significantly extend the network’s ability to resist collapse under amyloid stress.

**Figure 4.**
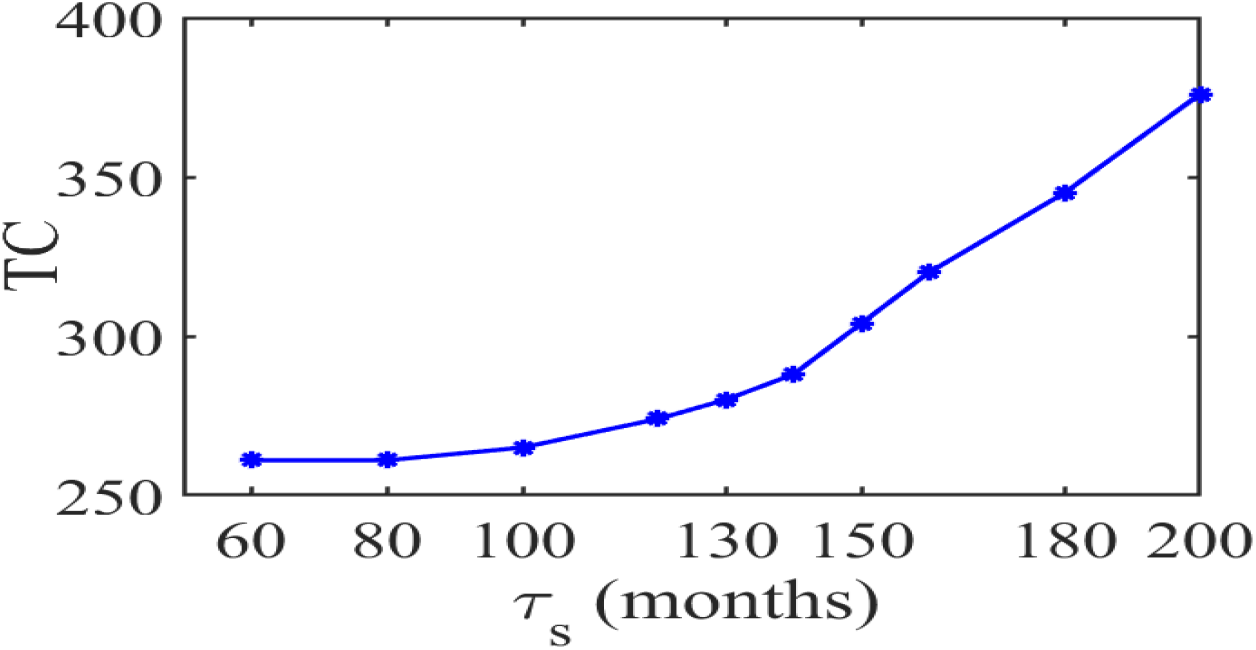
Critical collapse time increases with synaptic decay time: Relationship between *τ* _*s*_ and the critical collapse time *T*_*c*_. Longer-lasting synapses delay network failure, supporting the role of *τ* _*s*_ in compensatory resilience.

The approximately linear increase in *T*_*c*_ with *τ* _*s*_ supports the hypothesis that synaptic longevity buffers the network against cumulative damage, allowing for a prolonged phase of functional compensation. This result not only reinforces the biphasic degradation trajectory observed in Figure 3, but also suggests that *τ* _*s*_ serves as a key parameter in modulating the onset of catastrophic network failure. The clinical implication is that interindividual differences in synaptic decay dynamics may partially account for the observed variability in disease progression rates among AD patients [2,30].

From a theoretical standpoint, the nearly linear relationship between *τ* _*s*_ and *t*_*c*_ aligns with the idea of delayed transitions in complex systems, where slower decay rates permit extended stability until a critical threshold is breached [28]. This finding also highlights the potential of *τ* _*s*_ as a surrogate biomarker or a target for therapeutic modulation, suggesting that interventions aimed at slowing synaptic decay could meaningfully postpone the functional collapse of large-scale brain networks.

### 3.4 Effects of Rewiring Probability (prewire)

To investigate how network topology influences vulnerability to amyloid-induced collapse, we systematically varied the rewiring probability *p*_rewire_ in the Watts-Strogatz model, transitioning from ordered lattices ( *p* = 0 ) to small-world ( *p* = 0.1,0 .5 ) and fully random networks ( *p* = 1 ). Figure 5 illustrates the impact of topology on both the fraction of the Largest Strongly Connected Component (LSCC) and global efficiency (GE) over time. Ordered networks degraded rapidly following the crossing of the toxicity threshold ( *A*_th_ = 25 *CL* ), with LSCC collapsing shortly after *A* ≈ 50 CL. In contrast, small-world networks exhibited markedly enhanced resilience, maintaining both LSCC integrity and functional efficiency until amyloid levels approached high threshold ( ≈ 75 CL). Random networks showed intermediate behavior, where the redundancy of long-range connections delayed structural collapse, but we observed acceleration of the decline in GE due to loss of clustering.

**Figure 5.**
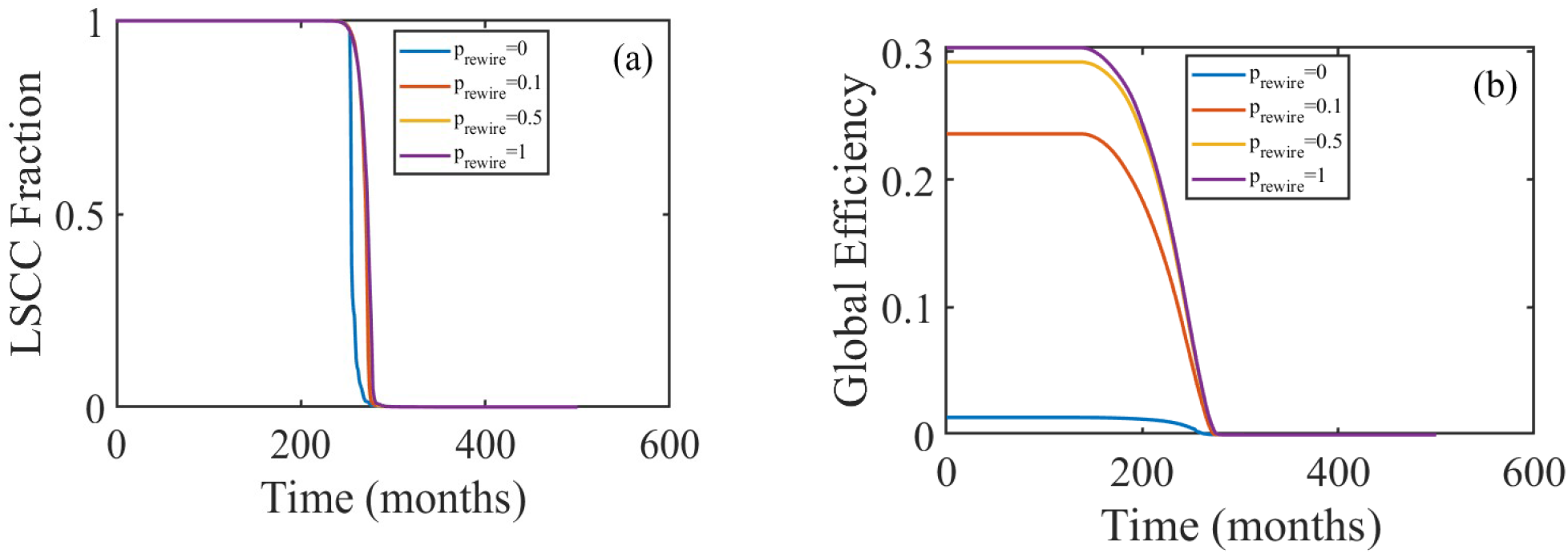
Network topology influences vulnerability to collapse: LSCC Fraction (a) and Global efficiency (b) over time for different rewiring probabilities *p*_rewire_ - Small-world topologies ( *p* = 0.1,0 .5 ) delay structural and functional collapse compared to ordered or random networks.

These results highlight a key structural insight: small-world topologies optimally balance local clustering and short path lengths, allowing them to absorb localized damage while maintaining global communication [32]. The delayed collapse observed in small-world networks provides a mechanistic explanation for the clinical dissociation between amyloid accumulation and the onset of cognitive symptoms [3], reinforcing the hypothesis that brain architecture has evolved for both efficiency and robustness. Furthermore, the finding that global efficiency consistently declines before LSCC failure across all topologies suggests that GE is a more sensitive and generalizable early-warning indicator of network failure. This aligns with the concept of critical slowing down as networks approach a tipping point [28]. Collectively, these results emphasize the importance of topological features in determining the timing and severity of AD-related network degradation, and suggest that individual differences in brain connectivity architecture could partly explain the heterogeneity of Alzheimer’s progression [5].

While the above results characterize the structural disintegration of the network, the functional consequences for spiking activity remain to be quantified. We next analyze how amyloid-induced degradation alters dynamical observables across disease stages

## 4 Functional Collapse of Spiking Dynamics

To characterize the functional consequences of amyloid-induced structural degradation, we simulated spiking dynamics across six key stages of disease progression ( t = 100,200,260,275,300,350 months) using an excitatory Izhikevich network augmented with short-term plasticity (STP). Neurons were simulated on static adjacency matrices extracted at each snapshot, reflecting cumulative synaptic loss due to amyloid burden. Isolated neurons (zero in-degree) were retained in the network to preserve effective network size but remained quiescent, consistent with biological constraints.

We synthesize multiple dynamical observables that reveal a nuanced, three-phase collapse. We observe the emergence of functional subpopulations. Dual-order raster plots sorting neurons by (i) total spike count and (ii) in-degree, uncover the progressive formation of three distinct subpopulations (Fig.6). In early stages ( ≤ 200 months), activity propagates as wave-like cascades from high-connectivity neurons to downstream units. By mid-stage ( *t* ≈ 275 months), a tripartite structure dominates: (1) a shrinking core (top 15% by spikes) exhibiting burst-like firing; (2) a large transient population firing sporadically; and (3) a growing silent fraction (bottom 30 % ) with no activity. In late stages ( ≥ 300 months), the core vanishes, the transient group fragments, and silence prevails ( > 70 % inactive).

The instantaneous population firing rate, 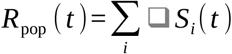, reveals a progressive degradation of collective dynamics (Fig7). In early stages ( ≤ 200 months), *R*_pop_ ( *t* ) exhibits robust, recurrent transients indicative of wave-like propagation and network excitability. By *t* = 275 months, these transients become sparse, irregular, and lower in amplitude, reflecting the fragmentation of functional modules. In late stages ( ≥ 300 months), *R*_pop_ ( *t* ) collapses to near-zero with only rare, uncorrelated spikes, confirming global hypoactivity.

As shown in Fig. 8a, the Fano factor increases dramatically, from ∼ 0.1 at *t* = 100 months to > 4 at *t* = 350 months-reflecting a transition from network-coordinated spiking to bursty, stochastic, and asynchronous firing. In stark contrast, the CV remains nearly constant across all disease stages (Fig. 8b), indicating Figure 8: Spiking irregularity during disease progression. (a) The coefficient of variation (CV) of interspike intervals remains approximately constant across all stages, indicating preserved intrinsic spiking regularity. (b) The Fano factor (variance-to-mean ratio of spike counts in 500 - *ms* sliding windows) increases from ∼ 0.1 to > 4, signaling a shift from coordinated network activity to pathological asynchrony and burstiness. Simulations use an excitatory Izhikevich network with STP at six disease stages (t=100 to 350 months) that intrinsic neuronal spiking regularity is preserved despite progressive synaptic loss. This dissociation demonstrates that functional collapse arises from the breakdown of network-level coordination, not from altered single-neuron dynamics.

In Fig.9, we observe that as disease progresses, the network undergoes a transition from synchronized, high activity states to desynchronized, low-activity regimes. While mean firing rate and global synchrony collapse after *t* = 275 months, the spectral profile retains its dominant gamma-band peaks; albeit with reduced power, suggesting that the mechanism for rhythm generation persists even as amplitude fades. This decoupling of frequency stability from power decline may reflect early-stage compensation followed by late-stage network failure.

Collectively, these results refine the original two-phase model into a more biologically grounded trajectory:

- A decoupled propagation phase ( ≤ 250 months): intact structure but disordered function, with emergent subpopulations, minimal hub dominance, and intact population transients.
- A reorganization phase (250 ≤ *t* ≤ 300 months): rising variability, spectral slowing, fragmented population bursts, and partial restoration of structure-function alignment as resilient hubs sustain residual activity.
- A collapsed silence phase ( ≈ 300 months): asynchrony, near-zero population rate, artifactual rank correlation, and loss of all collective dynamics.

This progression aligns with multimodal clinical data particularly the paradoxical coexistence of focal hyperactivity and widespread hypo-activity in early AD and establishes structured raster analysis, rank dynamics, and population rate decomposition as sensitive, multiscale biomarkers of network degradation.

Additionally, the inclusion of short-term plasticity (STP) in the model, particularly synaptic facilitation, may contribute to the maintenance of synchrony in early stages. STP dynamically enhances synaptic transmission during repetitive activity, thereby reinforcing transient coordination across neurons even as structural integrity degrades. While not isolated in this study, its role likely complements topological resilience in delaying functional disintegration [33].

## 5 Discussion

Our findings highlight the multifactorial nature of Alzheimer’s disease (AD) progression, in which molecular, structural, and dynamical elements interact across scales. Using a computational model that integrates amyloid accumulation, synaptic pruning, and biologically realistic neuronal dynamics, we revealed key mechanisms governing the resilience and collapse of brain networks in AD. First, our results reinforce the critical role of network topology. Small-world architectures, which balance high clustering with short path lengths, confer significant resilience against amyloid-induced degradation. These networks exhibit delayed collapse of both structural (LSCC) and functional (global efficiency) markers compared to ordered or random configurations (Figure 5), supporting the hypothesis that small-world connectivity has evolved to optimize robustness [32,34]. Notably, global efficiency declines prior to LSCC collapse across all topologies, suggesting that functional metrics may serve as early-warning indicators of network failure [28].

Second, we observed a clear dissociation between amyloid deposition and the timing of network collapse. The functional breakdown lags behind the molecular pathology (Fig. 1), mirroring clinical observations where amyloid accumulation precedes cognitive decline by a decade or more [2,3]. The inclusion of synaptic time constants ( *τ* _*s*_ ) in our model further refines this timeline: longer *τ* values extend the network’s compensatory phase, delaying collapse and increasing the critical amyloid burden required to trigger failure (Figures 3, 4). This aligns with the cognitive reserve hypothesis [30], suggesting that individuals with slower synaptic degradation may sustain function longer despite equivalent amyloid load.

Our spiking-network simulations reveal how amyloid-driven synaptic loss reshapes neuronal population dynamics across disease progression. By analyzing activity across six stages from preclinical ( = 100 months) to advanced dementia ( *t* = 350 months), we identify a three-phase functional trajectory that refines the traditional view of AD as a simple hypo-activity syndrome. First, dual-order raster plots (Fig. 6) uncover the progressive emergence of three functionally distinct subpopulations: (1) a hyperactive core (top 15% by spike count) exhibiting burst-like firing; (2) a large transient group with sporadic activity; and (3) a silent cohort (>70% by late stage) with no spiking output. This tripartite organization mirrors clinical electrophysiological data showing coexisting focal hyperactivity and widespread hypo-activity in early AD [10]. The alignment between structural connectivity (in-degree) and functional output (spike count) degrades over time, reflecting a progressive decoupling of structure and function. Second, the instantaneous population firing rate (Fig. 7) demonstrates a clear transition from coordinated, wave-like transients in early stages to fragmented, sparse bursts by mid-stage, and finally to near-complete silence in late stages. This confirms that network-level coordination not just mean firing rate is the first casualty of synaptic degradation. Third, the dissociation between Fano factor and coefficient of variation (CV) (Fig. 8) provides critical mechanistic insight. The Fano factor which measures spike-count variability across time; increases > 40 and fold (from ∼ 0.1 to > 4 ), signaling a shift from reliable, network-driven spiking to pathological burstiness and asynchrony. In stark contrast, the CV of interspike intervals remains stable, indicating that intrinsic neuronal spiking regularity is preserved throughout disease progression. This confirms that functional collapse arises from loss of network level coordination, not altered single-neuron dynamics, a key distinction for biomarker development and therapeutic targeting [10,35]. Finally, metrics of population synchrony and mean firing rate (Fig. 9) quantify the collapse of collective dynamics. Both decline sharply after amyloid surpasses ∼ 75 CL, coinciding with the disappearance of the hyperactive core and the fragmentation of the transient population. Together, these results establish that AD-related network failure is not uniform, but instead involves dynamic reorganization into functionally segregated subpopulations, followed by progressive disintegration of all coordinated activity.

**Figure 6.**
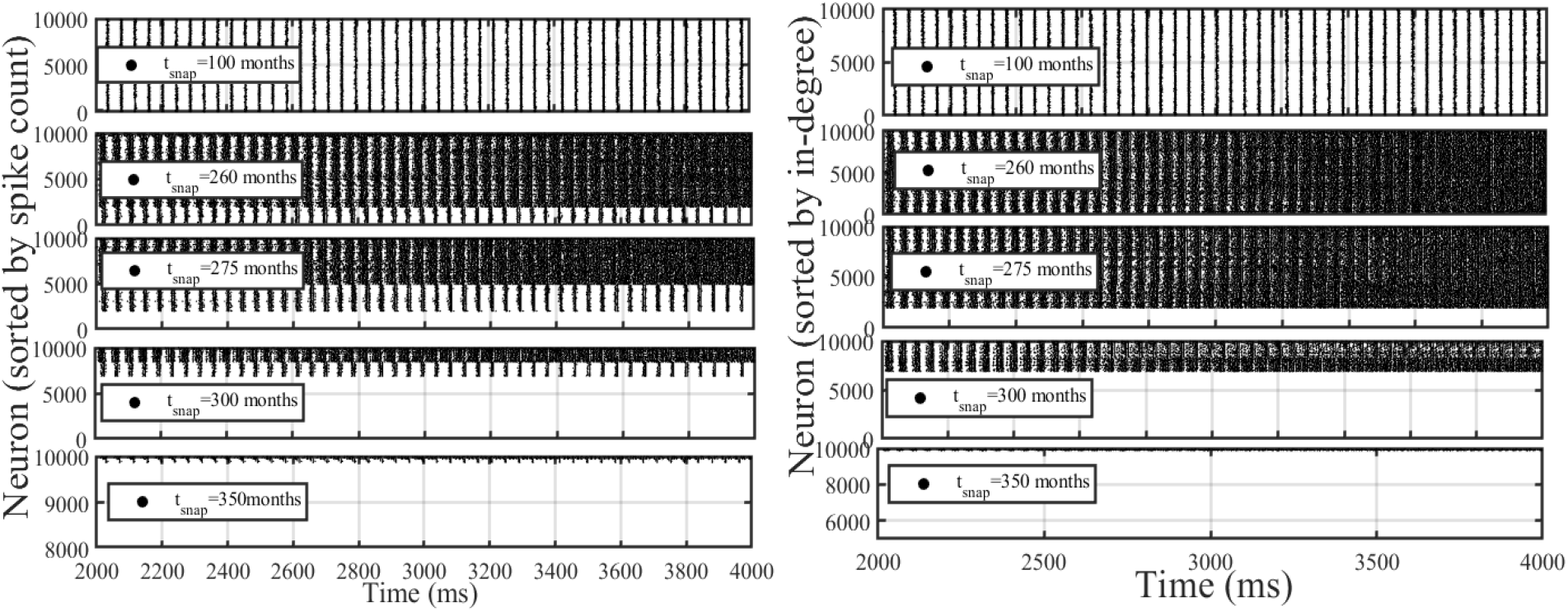
Dual-order raster plots (sorted by spike count and in-degree) reveal three emergent functional subpopulations: a hyperactive core, a transient group, and a silent cohort.

**Figure 7.**
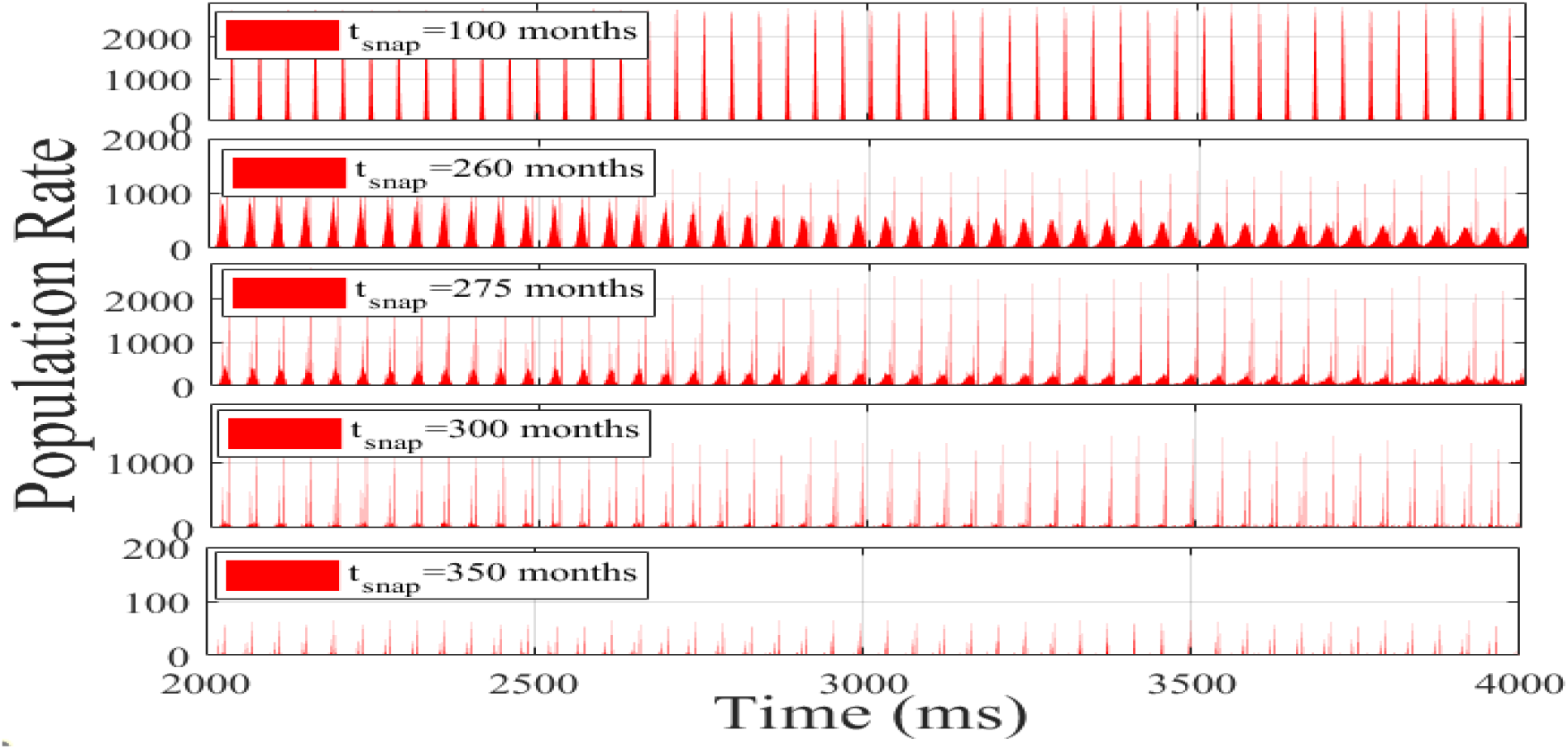
Instantaneous population firing rate showing loss of coordinated transients by mid-stage and near-silence in late stages.

**Figure 8.**
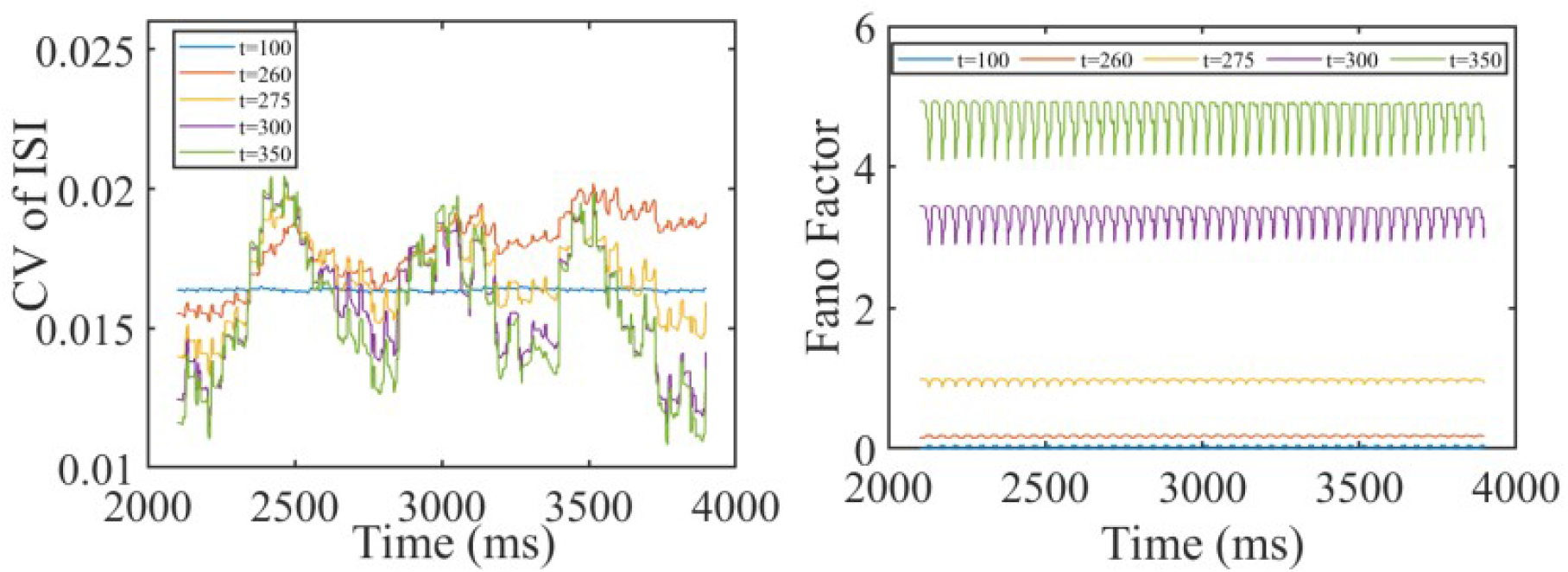
Spiking irregularity was quantified using two complementary metrics: the coefficient of variation (CV) of interspike intervals and the Fano factor (spike-count variance-to-mean ratio in 500 - *ms* sliding windows).

**Figure 9.**
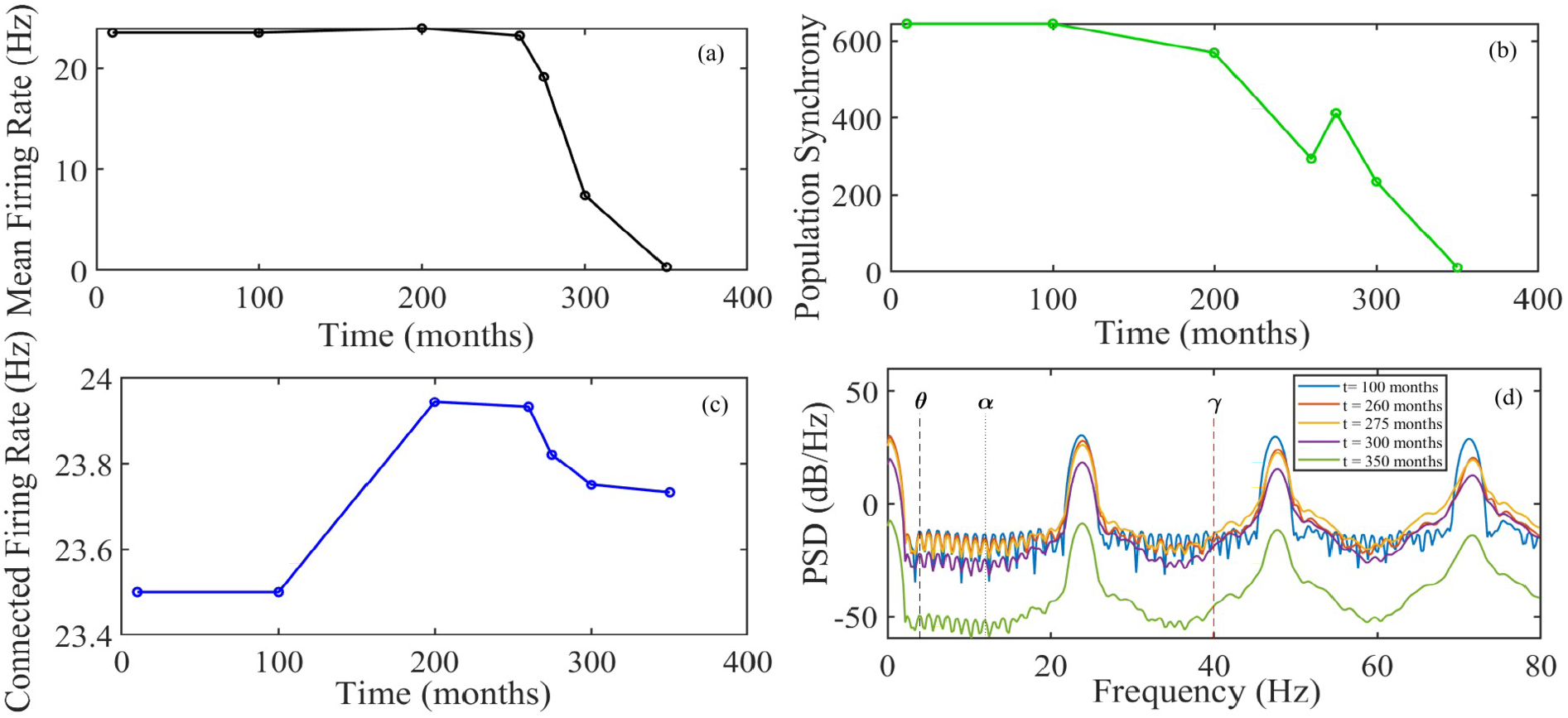
(a) Mean population firing rate declines sharply after *t* = 275 months. (b) Population synchrony (measured as global coherence or order parameter) decreases monotonically with disease progression.(c) Firing rate of connected neurons remains stable until *t* = 200, then drops slightly - suggesting preserved local activity despite global degradation. (d) Power spectral density (PSD) reveals persistent gamma-band oscillations ( 25 - 75 *Hz* ) across all stages, but power diminishes significantly at late stages ( *t* = 350 ), indicating loss of high-frequency synchrony. Theta ( *θ*,4 *Hz* ), alpha ( *α*,12 *Hz* ), and gamma ( *γ*,40 *Hz* ) bands are marked for reference. Parameters: *a* = 0.1, *b* = 0.2, *c* =-65, *d* = 0.2, *τ* _facil_ = 100 *ms, τ* _rec_ = 10 *ms, U* = 0.5.

These findings highlight the value of multiscale spiking, particularly structured raster analysis, Fano factor, and population synchrony, as sensitive biomarkers of early network dysfunction. They also reinforce the view of AD as a systems-level cascade, where synaptic loss disrupts emergent population dynamics long before global silence sets in.

Lastly, we note that the inclusion of short-term synaptic plasticity (STP), particularly facilitation, likely contributes to transient synchrony preservation during early disease stages. By dynamically enhancing synaptic efficacy during repeated activation, STP helps maintain coordinated spiking despite gradual structural decay. While not isolated in this study, its presence adds biological realism and plausibility to observed compensatory dynamics [33].

Although our model omits inhibitory neurons, our results align with broader principles observed in pathological network dynamics: namely, that the interplay between structural connectivity and activity propagation determines whether a perturbation dissipates or cascades. For example, in epilepsy, seizure-like inputs can either be contained or amplified depending on inhibitory control and network architecture [36]. In contrast, in amyloid-driven AD models, the gradual erosion of excitatory connectivity leads not to runaway excitation, but to a failure of propagation and eventual silencing, illustrating two divergent failure modes of brain network resilience.

In sum, our work emphasizes that Alzheimer’s disease cannot be fully understood without considering the interplay between intrinsic neuronal properties, synaptic dynamics, and global network topology. These insights support the use of multiscale computational models to identify early biomarkers, understand individual variability, and predict the impact of potential interventions. Future work should focus on integrating patient specific connectomes and cell-type heterogeneity to move toward personalized modeling and therapeutic targeting.

## 6 Conclusion

This study presents a multiscale computational framework that bridges molecular pathology and network-level dysfunction in Alzheimer’s disease. By modeling amyloid accumulation, synaptic degradation, short-term plasticity, and spiking neuron dynamics on biologically inspired small-world networks, we identified critical mechanisms governing the resilience and collapse of brain connectivity.

Theoretically, our findings demonstrate that small-world topologies confer superior resistance to amyloid induced failure, and that global efficiency serves as a robust early-warning indicator of collapse. Synaptic decay time constants ( *τ* _*s*_ ) modulate the delay between amyloid accumulation and functional breakdown, offering a biophysical explanation for the clinical dissociation between pathology and symptoms. Our spiking network simulations demonstrate that amyloid-induced synaptic loss drives a progressive and structured disintegration of population dynamics, culminating in global functional silence. Across six disease stages, we observe a three-phase trajectory: (i) an early propagation phase with wave-like, hub-driven activity; (ii) a mid-stage reorganization phase characterized by the emergence of three functional subpopulations, a hyperactive core, a transient group, and a growing silent cohort alongside rising population variability; and (iii) a late collapsed phase dominated by asynchrony, near-zero firing, and loss of all collective dynamics.

Critically, the Fano factor increases more than forty-fold while the coefficient of variation (CV) of interspike intervals remains stable, revealing a dissociation between population level desynchronization and preserved single-neuron spiking regularity. This confirms that network collapse arises from loss of connectivity not intrinsic neuronal dysfunction and establishes Fano factor, structured raster analysis, and population synchrony as sensitive, early biomarkers of network failure.

These findings provide a dynamic counterpart to structural models of Alzheimer’s progression and emphasize that functional degradation follows a non-uniform, hierarchical pattern long before global silence sets in. The framework offers a foundation for interpreting electrophysiological biomarkers in early AD and for evaluating interventions aimed at preserving network coordination.

Clinically, our results suggest that dynamic functional biomarkers, such as declining synchrony, rising entropy, and global efficiency fluctuations, may be more sensitive indicators of early dysfunction than static structural metrics. The model identifies the 25 CL threshold as the tipping point beyond which endogenous repair becomes ineffective, reinforcing the importance of early intervention. Delaying synaptic weakening or enhancing intrinsic plasticity could, in principle, extend network resilience by 50 - 100 months, depending on synaptic kinetics.

While our model integrates key biological and network-level features of Alzheimer’s progression, several limitations must be noted. First, the model focuses on amyloid-beta pathology and does not explicitly include tau protein dynamics, neuroinflammation, or glial interactions, which are known to influence disease progression. Second, we use homogeneous neurons and generic network topologies rather than patient specific connectomes or cell-type distributions. Third, while repair mechanisms and STP are included, other forms of plasticity such as long-term potentiation/depression are not modeled. Future work should extend this framework to incorporate multimodal pathology, heterogeneous connectome data, and personalized modeling to improve clinical translation.

Future work should focus on patient-specific modeling, incorporating individual connectomes, diverse neuron types, and co-pathologies such as tau propagation or neuroinflammation. The current framework provides a foundation for testing personalized interventions and for developing computational biomarkers to support early diagnosis and prognosis in Alzheimer’s disease. Ultimately, this approach offers both mechanistic insight and translational potential in the study of neurodegenerative network collapse.

## Abbreviations

AD: Alzheimer’s disease
*Aβ*: Amyloid-beta
CL: Amyloid concentration level
CV: Coefficient of variation
GE: Global efficiency
LSCC: Largest strongly connected component
PSD: Power spectral density
STP: Short-term plasticity

## Authors Contribution

ELFN: Conceptualization, Data Curation, Formal Analysis, Investigation, Methodology, Software, Validation, Visualization, Writing - Original Draft Preparation, Writing - Review & Editing.

DD: Conceptualization, Formal Analysis, Investigation, Methodology, Validation, Visualization, Supervision, Writing - Original Draft Preparation, Writing - Review & Editing.

FFF: Conceptualization, Formal Analysis, Funding Acquisition, Investigation, Methodology, Project Administration, Resources, Supervision, Validation, Writing - Original Draft Preparation, Writing - Review & Editing.

## Acknowledgment

FFF thanks the Brazilian National Council for Scientific and Technological Development (CNPq) and Sao Paulo Research Foundation (FAPESP). ELFN thanks Brazilian Federal Agency for Support and Evaluation of Graduate Education (CAPES). FFF and ELFN gratefully acknowledge the financial support by National Institute of Science and Technology in Innovative Research in Health Sciences – from Nanotechnology to Artificial Intelligence (INCT PICS) sponsored by Brazil’s National Council for Scientific and Technological Development (CNPq), grant no. 408417/2024-2, Coordination of Superior Level Staff Improvement (Capes), grant no. 88887.197686/2025-00, and São Paulo Research Foundation (FAPESP), grant no. 2025/26818-7.

## Data and Data coding

No empirical data generated. All code for model implementation, simulations, and figures is publicly available for reproducibility at: https://github.com/ediline000/amyloid-network-collapse.git

## Ethics statement

Not required – purely computational study (no human/animal subjects or primary biological data).

